# A biomimetic SPARC platform coupling adaptor recruitment and BRS-Lck condensation for enhanced NK/T cell combination therapy

**DOI:** 10.64898/2026.11.02.763574

**Authors:** Qi Wang, Songfang Wu, Yueting Qiu, Jizhong Lou, Changjie Lou, Hui Chen

## Abstract

Immune cell engineering has proven to be a promising therapeutic strategy for cancers and autoimmune diseases. Compared to classical CAR therapies, engineered T cells with modified TCRs (such as STAR-T and TRuC-T) exhibit lower tonic signaling and higher persistence. Inspired by this, we developed novel Synthetic Parallel Antigen Receptor Complexes (SPARC) based on the NK cell receptor 1 (NCR1) and the TCRα chain. These SPARCs can recognize target cells via engineered extracellular domains and transduce signals by recruiting endogenous adaptor proteins (CD3ζ/FcRγ). In NK cells, SPARCs confer high antigen specificity against lymphoma and enables NK cells to specifically recognize and kill tumor cells. Mechanistically, the cytoplasmic domain of NCR1 forms condensates with Lck kinase through two conserved basic-rich sequences (BRS), thereby promoting receptor clustering and enhancing NK cell cytotoxicity. Our findings indicate that SPARCs can also be applied to other immune cells expressing CD3ζ and FcRγ. Incorporation of costimulatory domains further improves the performance of SPARCs. Furthermore, combining SPARC-engineered T cells and NK cells showed superior efficacy compared to either cell type alone. This biomimetic design preserves low basal signaling, overcomes the recognition limitations of endogenous receptors, and provides a versatile platform for engineering other membrane complexes.

## Introduction

The engineering of immune cells has emerged as a transformative approach for cancer and autoimmune diseases, offering durable therapeutic responses. Chimeric antigen receptor (CAR) engineering, a representative strategy, has been widely applied to T cells and NK cells and achieved remarkable success in hematological malignancies. Nevertheless, CAR-associated issues such as tonic signaling, limited persistence, and antigen escape continue to drive next-generation immune cell engineering platforms.^1–3^ These challenges highlight the need for receptors that balance robust antitumor activity with clinical safety.

A central challenge is optimizing the balance between synthetic receptor sensitivity and safety. Overly robust immune activation can precipitate severe cytokine release syndrome (CRS) and neurotoxicity, whereas insufficient activation fails to eradicate tumors. Consequently, researchers seek an optimal equilibrium between sufficient sensitivity to recognize tumor cells with low antigen density and adequate safety to avert off-target effects.^4–6^ This equilibrium is particularly important because receptor affinity, signaling strength, and co-stimulation collectively determine the therapeutic window of engineered cells. Thus, rational receptor design must consider both antigen recognition and downstream signal quality.

TCR-T therapy is another promising adoptive cell immunotherapy, particularly for solid tumors.^7, 8^ By genetically modifying T cells to express tumor-specific TCRs, this approach engages the complete TCR signaling cascade, potentially increasing intracellular signaling complexity and reducing CRS risk.^9^ However, TCR-T is constrained by MHC restriction and potential alloreactivity.^10^ These limitations have motivated alternative architectures that combine antibody-type antigen recognition with TCR-derived signaling to broaden applicability while preserving natural signaling quality. Such designs may extend TCR-T beyond MHC-restricted antigens and improve safety.

Innovative strategies such as STAR-T and TRuC-T integrate antibody-derived recognition domains with TCR subunits, preserving native TCR signal transduction while expanding antigen recognition.^11–14^ These engineered cells show improved cytokine secretion, cytotoxicity, and persistence in vitro and in animal models. Such advances highlight that receptor architecture—extracellular conformation, transmembrane interactions, and intracellular signaling motifs—critically shapes activation thresholds, signal amplification, and effector duration.^15^ This understanding provides a theoretical foundation for rational design and motivates the search for natural signal-transduction templates that may overcome existing bottlenecks.^16^ It also suggests that modular receptor engineering can tune signaling strength without sacrificing specificity.

NK cells are promising effectors for cell-based immunotherapy, and CAR-NK is widely pursued.^17^ However, current CAR-NK platforms face limited persistence, tumor microenvironment-mediated suppression, and insufficient co-stimulation for full activation.^18, 19^ These limitations underscore the need for innovative engineering paradigms. NK receptors often mediate sensitive target recognition without constitutive activation. Similar to TCR, NK cell activating receptors often have short cytoplasmic tails lacking signaling motifs,^20^ and transduce activation signals through associated adaptors such as FcRγ, CD3ζ, and DAP12.^21–23^ Co-stimulation is also critical: first-generation CAR-NK cells relying solely on CD3ζ showed limited antitumor activity, and CD28 or 4-1BB domains have since been incorporated.^24–31^ Accumulating evidence indicates that costimulatory domains compatible with NK cell-intrinsic signaling are pivotal for maximizing antitumor efficacy and clinical translation.^32–34^ Therefore, selecting an appropriate costimulatory module is essential for next-generation CAR-NK development.

Here, we describe the development of Synthetic Parallel Antigen Receptor Complexes (SPARCs), a novel class of chimeric receptors that combine single-chain variable fragments (scFvs) as extracellular recognition domains with the signaling frameworks of NCR1, KIR3DS1, and the TCRα chain. These receptors form receptor complexes with endogenous CD3ζ, FcRγ, or DAP12 adaptor proteins at the cell surface, and initiate immune cell activation after specific antigen recognition. In NK cells, the NCR1-based SPARC confers high antigen specificity and enhanced cytotoxicity against lymphoma, mediated by the formation of NCR1-Lck signaling condensates. Addition of costimulatory domains to the SPARC further augmented the antitumor efficacy of NK cells. Moreover, we demonstrate that SPARCs are functionally transferable to other immune cell types, and that combined use of SPARC-engineered T and NK cells results in superior antitumor efficacy. This work establishes SPARCs as a versatile, low-basal-signaling platform for engineering immune receptor complexes with broad therapeutic potential.

## Results

### Design and validation of chimeric receptor complexes

The activating receptor complexes on NK cells include the NCR family members (such as NCR1 and NCR3), Fc receptors such as CD16 which mediates ADCC, and the short-tail members of the KIR family, KIR2DS and KIR3DS. NCR1^35^, NCR3^36, 37^, and CD16^38, 39^ associate with the adaptor proteins CD3ζ and FcRγ, whereas KIR2DS and KIR3DS bind to DAP12^40^ (**Figure 1A**). From these receptor families, we selected and designed synthetic receptors based on NCR1, NCR3, CD16, and KIR3DS1, all incorporating the single-chain antibody FMC63 (targeting CD19) as the extracellular recognition domain (**Figure 1B**).

**Figure 1.**
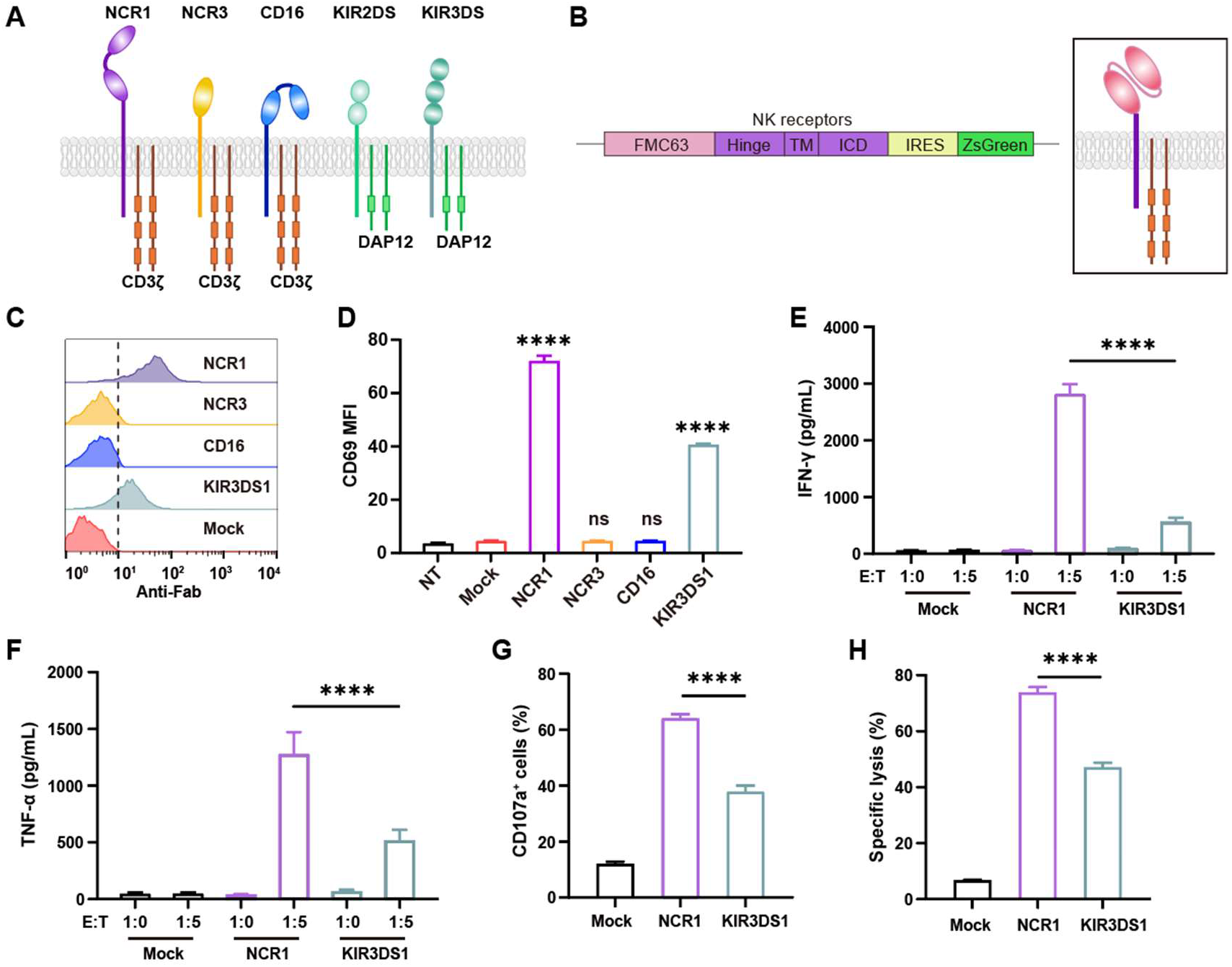
Design and validation of CD19-targeting chimeric receptor complexes derived from multiple NK cell activating receptors. **(A)** Schematics of NK cell activating receptors. **B** Gene (left) and protein (right) schematics of the chimeric receptors. **C** Membrane surface expression of chimeric receptors on NK92MI cell line determined by flow cytometry. **D** Mean fluorescence intensity (MFI) of CD69 expression on transduced NK92MI cells, determined by flow cytometry, after 24 hours coculture with CD19⁺ Nalm6 cells. (**E and F**) Cytokine productions of chimeric receptors engineered NK92MI cells upon Nalm6 cells stimulation for 24 hours. IFN-γ (**E**) and TNF-α (**F**) were measured. **G** Degranulation marker CD107a of chimeric receptors engineered NK92MI cells upon Nalm6 cells stimulation for 24 hours at an effector:target (E:T) ratio of 1:1. **H** Cytotoxicity of chimeric receptors engineered NK92MI cells upon Nalm6 cells stimulation for 24 hours at an effector:target (E:T) ratio of 1:1.Data are presented as mean with SD (n = 3; ∗∗∗∗p < 0.0001; ns, nonsignificant; unpaired two-tailed Student’s t-test for comparisons).

We transduced these chimeric receptors into the NK92MI cell line using lentiviral particles to assess their functionality. Surface expression was examined by antibody staining and flow cytometry. NCR1- and KIR3DS1-based receptors were efficiently displayed on the cell membrane, While NCR3- and CD16-based receptors showed no appreciable membrane expression (**Figure 1C and S1A**), this lack of expression may be attributable to their relatively short hinge between Ig-like domain and the transmembrane domain (hinge length: NCR1: 41aa; NCR3: 7aa; CD16: 13aa; KIR3DS1: 25aa).

Following stimulation with CD19-positive Nalm6 target cells, the four chimeric receptor-transduced NK cell populations exhibited differential activation, as reflected by the up-regulation of the early activation marker CD69 (**Figure 1D and S1B**). NK cells bearing NCR1- or KIR3DS1-based receptors were robustly activated, whereas those with NCR3- or CD16-based constructs failed to trigger NK cell activation due to impaired membrane expression.

### NCR1-based synthetic receptors elicit stronger NK cell activation than KIR3DS1-based receptors

After preliminary assessment of early NK cell activation signals, we next sought to further evaluate the function of SPARC-engineered NK cells. SPARC-engineered NK cells were co-cultured with Nalm6 target cells, and the secretion of IFN-γ and TNF-α was measured. We found that both constructs effectively activated NK cells and induced IFN-γ and TNF-α secretion, whereas the NCR1-based SPARC triggered significantly higher cytokine production than the KIR3DS1-based SPARC (**Figure 1E and 1F**). In addition, NK cells engineered with the NCR1-based SPARC exhibited substantially enhanced degranulation, as indicated by increased surface expression of CD107a, relative to those bearing the KIR3DS1-based SPARC (**Figure 1G**). Moreover, the NCR1-based SPARC conferred stronger cytotoxic activity against target cells (**Figure 1H**). The enhanced functionality of the NCR1-based SPARC may be attributable to its higher membrane expression (**Figure 1C and S1A**). Taken together, these results indicate that the NCR1-based SPARC effectively mediates NK cell activation and cytotoxicity against target cells.

### Activation of both NCR1- and KIR3DS1-based SPARCs requires transmembrane-mediated adaptor coupling

To verify that NK cell activation by SPARC constructs depends on their assembly with adaptor proteins at the cell membrane, we designed two kinds of variants in which the transmembrane domain containing the adaptor-binding site was either replaced with that of CD8a or mutated to uncharged alanine residues (**Figure 2A**). Flow cytometric analysis indicated that the NCR1-based SPARC exhibited enhanced cell-surface expression when its transmembrane domain failed to bind the adaptor protein (**Figure 2B**). Co-culture experiments demonstrated that variants lacking the adaptor-binding site exhibited significantly reduced NK cell activation, confirming that its activation is highly dependent on coupling with FcRγ (**Figure 2C**). Notably, even after losing the ability to bind adaptors, the NCR1-based SPARC retained partial activity.

**Figure 2.**
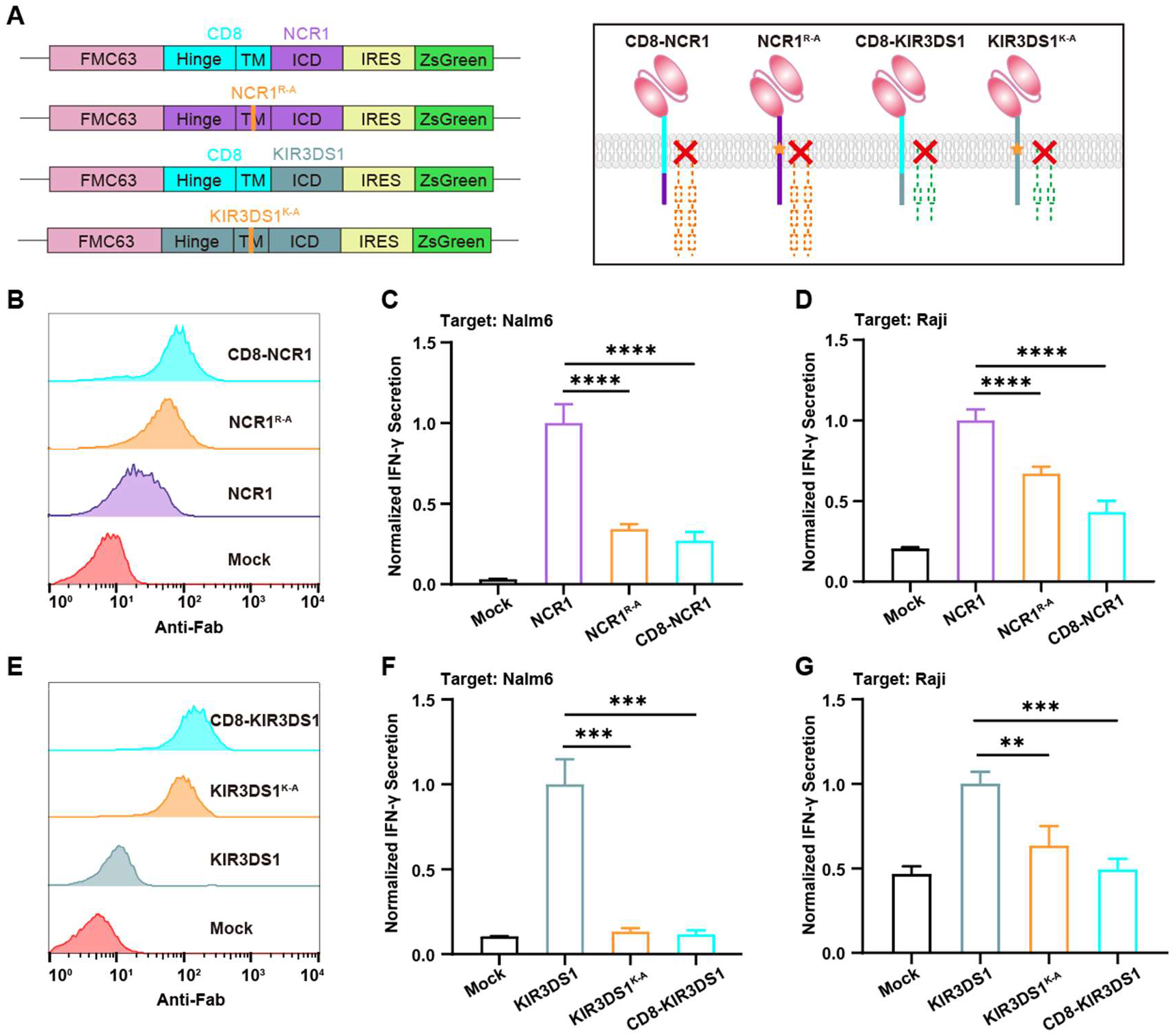
SPARC activation through NCR1 or KIR3DS1 relies on coupling to adaptor proteins via the transmembrane region. **A** Gene (left) and protein (right) schematics of the chimeric NCR1 and KIR3DS1 receptors with CD8α transmembrane domain or transmembrane domain mutation. **B** Membrane surface expression of chimeric NCR1 receptors on NK92MI cell line determined by flow cytometry. (**C and D**) Normalized IFN-γ secretion of chimeric NCR1 receptors engineered NK92MI cells upon Nalm6 **C** and Raji (**D**) cells stimulation for 24 hours at an effector:target (E:T) ratio of 1:5. **E** Membrane surface expression of chimeric KIR3DS1 receptors on NK92MI cell line determined by flow cytometry. (**F and G**) Normalized IFN-γ secretion of chimeric NCR1 receptors engineered NK92MI cells upon Nalm6 **F** and Raji (**G**) cells stimulation for 24 hours at an effector:target (E:T) ratio of 1:5. Data are presented as mean with SD (n = 3; ∗∗∗p < 0.001; ∗∗∗∗p < 0.0001; ns, nonsignificant; unpaired two-tailed Student’s t-test for comparisons).

To further exclude the influence of target cells, we used Raji cells as target cells in co-culture experiments and verified that this residual signal was target-cell independent and likely originated from the receptor itself (**Figure 2D**). We therefore introduced the corresponding transmembrane domain mutations and replacements into the KIR3DS1-based SPARC (**Figure 2A**). The resulting mutants were expressed normally on the cell membrane but failed to mediate cytokine secretion by NK cells (**Figure 2E-G**). These results suggest that, for the NCR1-based SPARC, in addition to adaptor-mediated signaling, the NCR1 intracellular domain may mediate NK cell activation through other mechanisms.

### Phase separation mediated by the NCR1 intracellular domain enhances transmembrane signal transduction of SPARC

Since the NCR1 intracellular domain is short and lacks any known motifs that directly transduce signals, the mechanism by which it promotes transmembrane signal transduction remains unclear. Cross-species conservation analysis of NCR1 revealed two highly conserved basic-rich sequence (BRS) regions within its cytoplasmic domain across diverse species (**Figure 3A**). These BRS regions could be implicated in phase separation, a mechanism critical for immune receptor signaling, as demonstrated in TCR and CD28.^41, 42^

**Figure 3.**
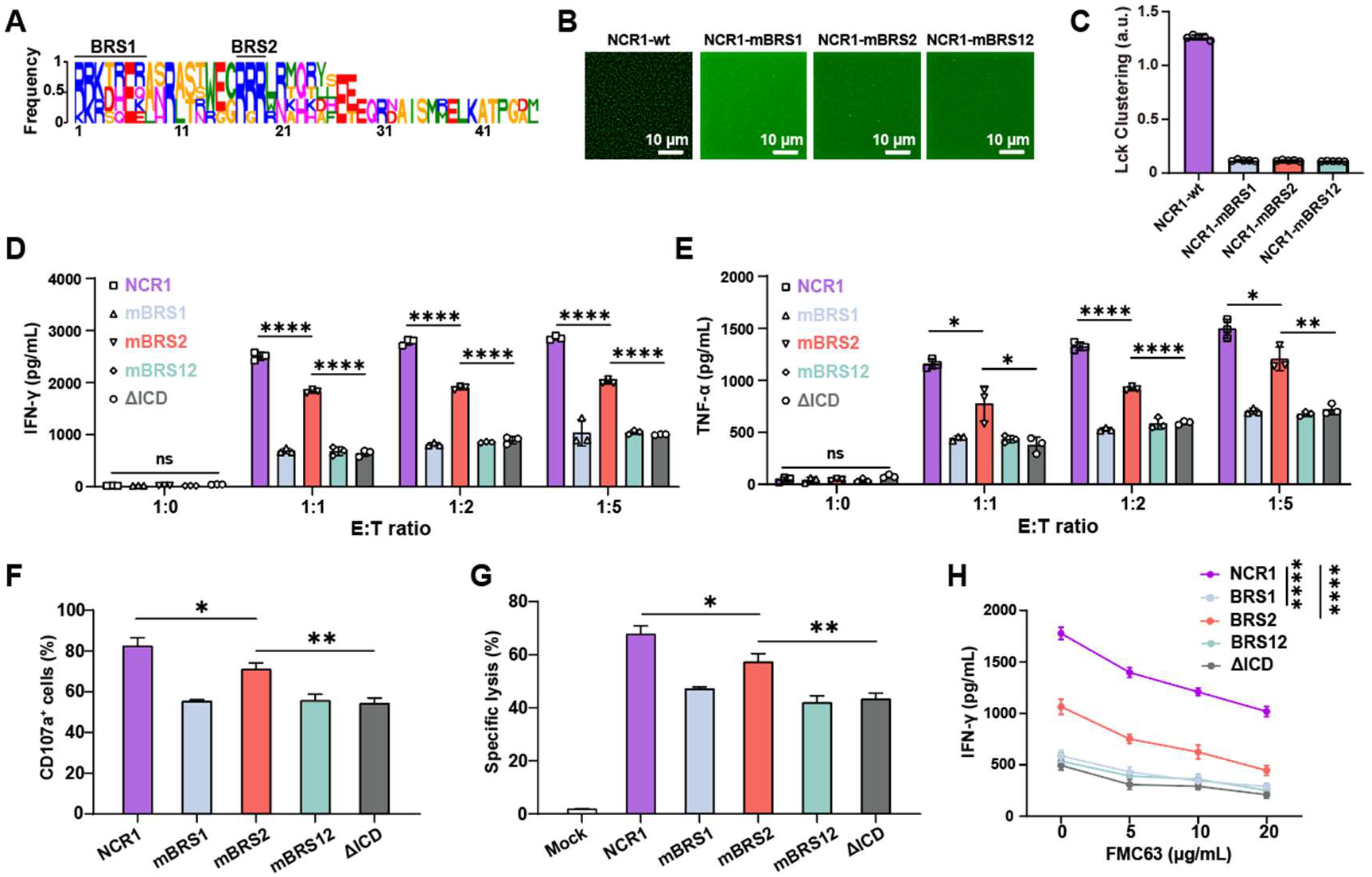
NCR1 intracellular domain enhances transmembrane signal transduction via phase separation. **A** Cross-species conservation analysis of NCR1 intracellular domain. **B** On-membrane phase separation between NCR1 intracellular domain mutations and Lck. Lck UD-SH3-SH2 was coated on supporting lipid bilayer (SLB), NCR1 cytoplasmic tail was added and incubated at room temperature for 30 min. **C** Lck clustering in images shown in (**B**) was quantified as described in Materials and Methods. (**D and E**), Cytokine productions of NCR1-based SPARC and its mutations engineered NK92MI cells upon Nalm6 cells stimulation for 24 hours at the indicated effector:target (E:T) ratios. IFN-γ (**D**) and TNF-α (**E**) were measured. **F** Degranulation marker CD107a of NCR1-based SPARC-NK92MI cells upon Nalm6 cells stimulation for 24 hours at an effector:target (E:T) ratio of 1:1. **G** Cytotoxicity of NCR1-based SPARC-NK92MI cells upon Nalm6 cells stimulation for 24 hours at an effector:target (E:T) ratio of 1:1. **H** IFN-γ secretion of NCR1-based SPARC-NK92MI cells upon Nalm6 cells blocked by soluble FMC63 stimulation for 24 hours at an effector:target (E:T) ratio of 1:1. Data are presented as mean with SD (n = 3; ∗, P < 0.05; ∗∗, P < 0.01; ∗∗∗∗p < 0.0001; ns, nonsignificant; unpaired two-tailed Student’s t-test for comparisons).

Using an in vitro artificial planar membrane system, we observed that the NCR1 cytoplasmic domain exhibits phase separation capability with Lck kinase, a key phosphorylating enzyme in NK cell activation (**Figure 3B**). The phase separation activity was strictly dependent on the two BRS regions: mutating positively charged residues in either BRS to neutral alanine abolished this capability (**Figure 3C and Figure S2A**). This finding suggests that the NCR1 cytoplasmic domain may recruit endogenous receptors (e.g., native NCR1) and kinases (e.g., Lck) via phase separation in vivo, thereby locally enriching receptors and signaling molecules at the NK cell membrane to amplify immune activation.^43^ This mechanism provides a plausible explanation for the low-level antigen-specific activation observed in the absence of adaptor protein binding.

To further evaluate the functional impact of BRS-mediated phase separation, we engineered SPARCs with BRS mutations. Flow cytometry characterization shows that the expression of various SPARCs on the membrane is basically consistent (**Figure S2B**). To directly visualize receptor clustering during activation, we performed TIRF (total internal reflection fluorescence) microscopy on NK cells expressing fluorescently tagged SPARCs (**Figure S2C**). When seeded onto CD19-coated slides, SPARC containing the NCR1 cytoplasmic domain exhibited pronounced membrane clustering at antigen contact sites. In contrast, SPARCs with BRS mutations or lacking the cytoplasmic domain showed less clustering, confirming that phase separation mediated by the NCR1 cytoplasmic domain enhances SPARC clustering (**Figure S2D and S2E**).

Disruption of either BRS motif significantly reduced antigen-specific NK cell activation, as evidenced by diminished cytokines secretion (**Figure 3D and 3E**). In addition, BRS mutation led to partially impaired degranulation, as indicated by CD107a surface expression (**Figure 3F**), and reduced target cell lysis efficiency (**Figure 3G**). These results suggest that BRS mutations affect SPARC clustering on NK cells and thereby impair cytokine secretion by NK cells. Receptor activation is an avidity-governed process, therefore, BRS-enhanced SPARC clustering may confer greater sensitivity on SPARC. To examine the responsiveness of SPARCs to low-density antigens, we titrated graded concentrations of FMC63 (the single-chain antibody used as the antigen-binding moiety in the chimeric receptors) to block CD19 on Raji cells, thereby simulating a range of target antigen densities, and subsequently co-cultured the SPARC-NK cells with these target cells. IFN-γ secretion from NK cells revealed that the intact NCR1 cytoplasmic domain confers superior antigen-specific activation and maintains robust sensitivity even under low-antigen-density conditions (**Figure 3H**). Notably, this sensitivity was strictly dependent on the two BRS motifs within the intracellular domain. Collectively, these results underscore the indispensable role of BRS-driven phase separation in amplifying immune responses.

Taken together, we found that NCR1-based SPARC mediates NK cell activation through two mechanisms: a canonical adaptor-dependent signaling branch, and a phase separation-dependent branch in which the BRS motif coalesces with Lck kinase and thereby licenses engagement of endogenous NK receptors for downstream cascade activation.^43^ When both the adaptor-binding sites in the transmembrane domain and the BRS motifs in the intracellular domain were simultaneously mutated, the NCR1-based SPARC could no longer be specifically activated by target cells (**Figure S2F**).

### SPARC represent a promising engineering platform for T cell-based immunotherapy

We next sought to extend the design of our chimeric receptor complexes. In the native T cell receptor (TCR) complex, the TCRα chain possesses a transmembrane domain that can associate with the adaptor protein CD3ζ to assemble functional receptor complexes.^44^ Since the TCRα chain contains a positively charged lysine (K) residue that binds the CD3δε dimer via electrostatic interactions, we mutated K to the uncharged residues alanine (A) or isoleucine (I) and used these variants as the transmembrane domain of SPARC. We incorporated the TCRα transmembrane domain mutants into our SPARC constructs (**Figure 4A**) and observed that these SPARCs were successfully expressed on the surface of NK cells (**Figure 4B**) and could mediate NK cell cytokine secretion in a ligand dependent manner (**Figure 4C**).

**Figure 4.**
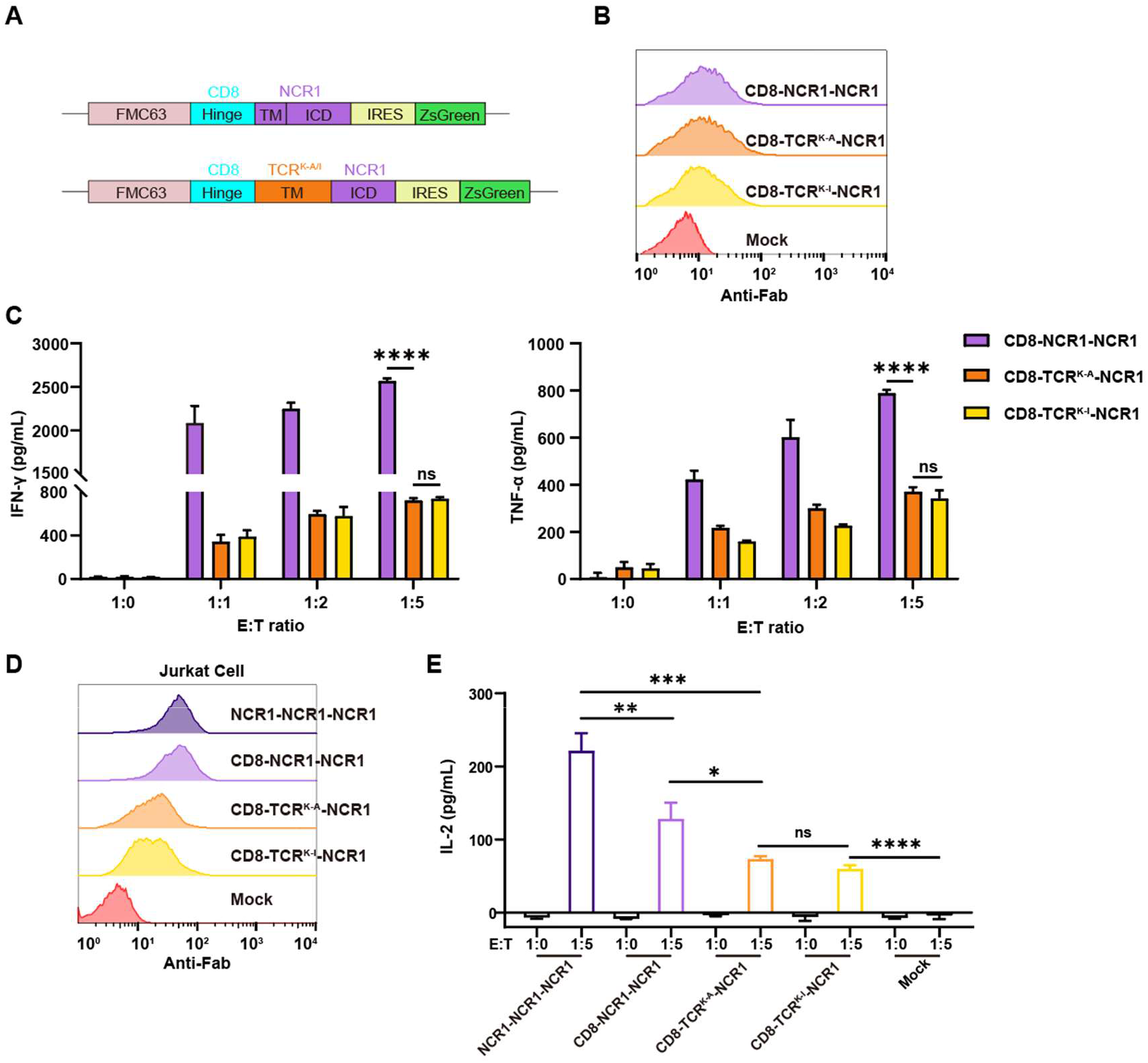
TCR-derived transmembrane domains support functional SPARC in both NK and T cells. **A** Gene schematics of the chimeric NCR1 receptors with CD8α stalk and TCRα mutation transmembrane domain. **B** Surface expression of SPARCs in (**A**) on NK92MI cells by flow cytometry. **C** Cytokine productions of SPARC-NK92MI cells upon Nalm6 cells stimulation for 24 hours at the indicated effector:target (E:T) ratios. IFN-γ (left) and TNF-α (right) were measured. **D** Surface expression of SPARC-Jurkat cells by flow cytometry. **E** IL-2 secretion of SPARC-Jurkat cells upon Raji cells stimulation for 24 hours. Data are presented as mean with SD (n = 3; ∗, P < 0.05; ∗∗, P < 0.01; ∗∗∗p < 0.001; ∗∗∗∗p < 0.0001; ns, nonsignificant; unpaired two-tailed Student’s t-test for comparisons).

Similar to NK cells, T cells are capable of tumor cell killing and represent the most commonly used engineered vehicles for targeted cancer immunotherapy. Given that CD3ζ is also present in T cells as a critical adaptor, our SPARCs hold considerable promise for T cell engineering. To explore this possibility, we tested SPARCs in the Jurkat T cell line and found that SPARCs bearing either the NCR1 or the TCRα transmembrane domain could be expressed on Jurkat cell surface (**Figure 4D**) and specifically activate Jurkat cells, as measured by IL-2 secretion (**Figure 4E**). Both TCRα mutants exhibited comparable activation levels, but these were weaker than those conferred by the NCR1 transmembrane domain. This may be because SPARC containing the TCRα transmembrane domain shows lower membrane expression than SPARC containing the NCR1 transmembrane domain (**Figure 4D**). Collectively, these findings indicate that SPARC is a promising platform for both NK cell and T cell-based applications.

### SPARC-based T-cell and NK-cell combination therapy

Conventional CAR designs often incorporate costimulatory receptor ICD to provide more robust intracellular signals, thereby enhancing immune cell activation and persistence. We likewise sought to optimize SPARC-NK function by incorporating signaling from the costimulatory receptor 4-1BB into SPARC (**Figure 5A**). Flow cytometry showed that the surface expression level of the second-generation CAR was higher than that of the SPARC (**Figure 5B**). Upon target cell stimulation, SPARC bearing the costimulatory ICD exhibited comparable or superior activation levels, as measured by the secretion of IFN-γ and TNF-α (**Figure 5C**).

**Figure 5.**
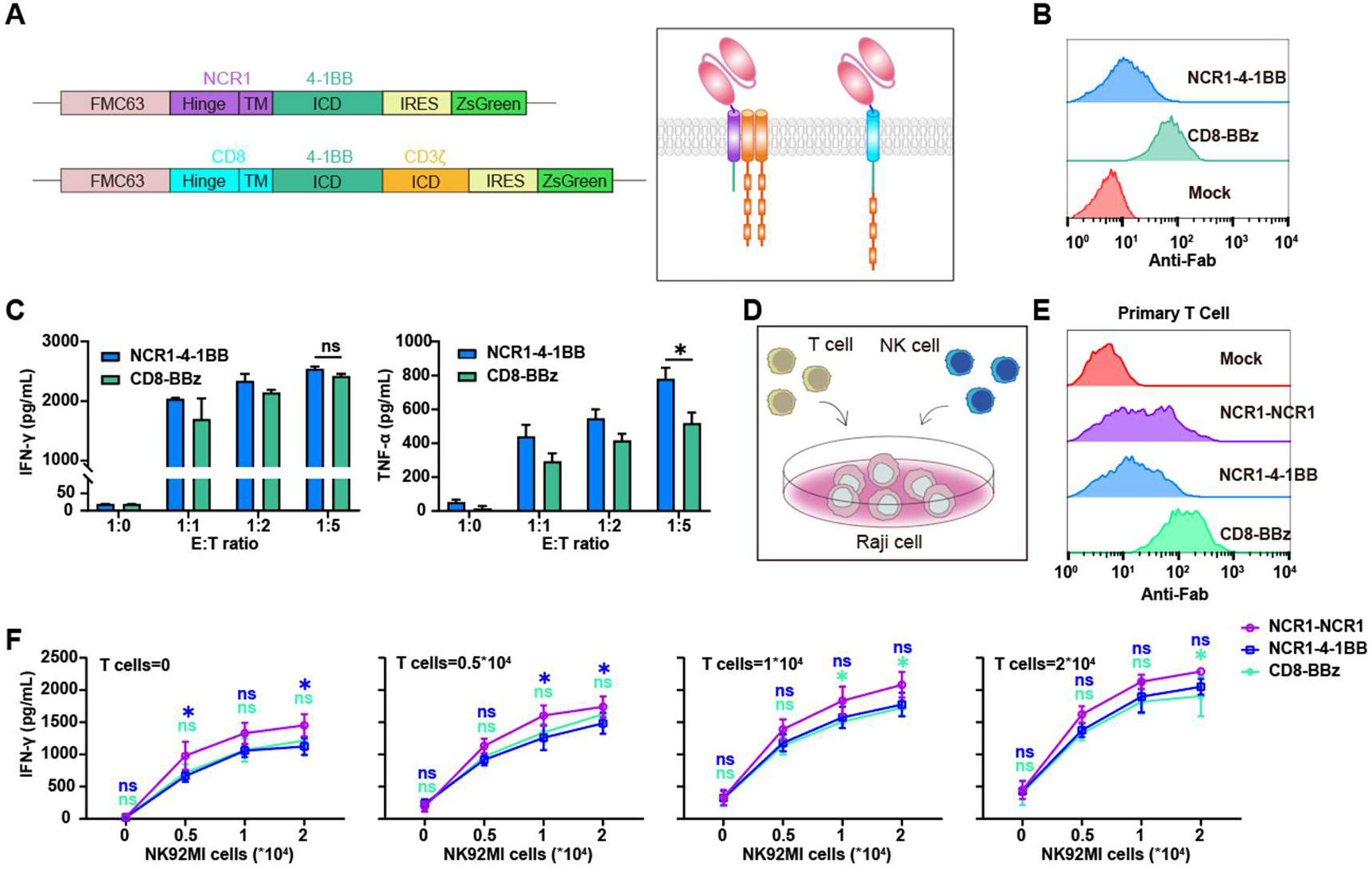
Incorporating the 4-1BB co-stimulatory domain into SPARC enables SPARC-based NK and T cell combination therapy to achieve efficacy equivalent to that of conventional second-generation CAR. **A** Gene (left) and protein (right) schematics of the SPARC with 4-1BB co-stimulatory domain and BBz CAR. **B** Surface expression of SPARC and CAR on NK92MI cells by flow cytometry. **C** Cytokine productions of SPARC-NK92MI cells upon Nalm6 cells stimulation for 24 hours at the indicated effector:target (E:T) ratios. IFN-γ (left) and TNF-α (right) were measured. **D** Schematic representation of the in vitro experiment for primary T cells and NK92MI cells combined therapy. **E** Surface expression of SPARCs and CAR on primary T cells by flow cytometry. **F** IFN-γ secretion in the supernatants from co-culture of SPARC- and CAR-NK92MI cells and T cells upon Raji cells stimulation for 24 hours at the indicated cell numbers. Raji cells is 20000 per well. Data are presented as mean with SD (n = 3; ∗, P < 0.05; ns, nonsignificant; unpaired two-tailed Student’s t-test for comparisons).

We additionally generated chimeric receptor constructs incorporating the CD28 intracellular domain and compared their properties with those of a second-generation CAR 28z (**Figure S3A**). Flow cytometric analysis revealed that SPARC composed of the NCR1 transmembrane domain and the CD28 cytoplasmic domain exhibited undetectable surface expression (**Figure S3B**). Upon stimulation with target cells, NK cells engineered with this NCR1-CD28 SPARC showed only modest activation, which was markedly lower than that observed for 28z and BBz (**Figure S3C**). The poor surface expression of NCR1-CD28 SPARC may be attributed to rapid internalization or degradation following membrane trafficking, whereas the residual activation induced by target cell stimulation likely results from engagement of the small fraction of surface-expressed receptors with target antigens.

Given the frequent crosstalk between NK and T cells in the immune system, which mutually regulate immune functions while both possessing tumor-killing capacity, previous studies have suggested potential synergism between NK and T cells in tumor clearance.^45–47^ Moreover, cell-based therapies often face challenges of tumor heterogeneity, and combined treatment with NK and T cells may better potentiate immune responses and tumor eradication.^48–51^ Since we have demonstrated that our SPARC is applicable to T cells, we sought to exploit the synergistic effects of both cell types by engineering them with the same SPARC and administering them in combination.

To test this, we performed in vitro co-culture experiments using titrated mixtures of engineered T cells and NK cells alongside CD19-expressing Raji lymphoma cells (**Figure 5D**). SPARC has lower expression levels on the surface of T cells compared to BBz CAR (**Figure 5E**). Strikingly, the combined therapy demonstrated superior cytokine secretion compared to monotherapy with either cell type alone (**Figure 5F**). Notably, SPARC retaining the NCR1 cytoplasmic domain outperformed SPARC with 41BB cytoplasmic domain or BBz CAR in promoting synergistic activation, as evidenced by enhanced cytokine production (**Figure 5F**).

The superior efficacy of combined therapy likely arises from bidirectional crosstalk: NK cells amplify T cell activation through cytokine secretion (e.g., IL-12, IL-15), while T cells enhance NK cell cytotoxicity via membrane-bound ligands (e.g., CD40L).^52^ Additionally, the NCR1 cytoplasmic domain’s phase separation capability may facilitate the formation of spatially organized signaling hubs at immune synapse interfaces, further amplifying cooperative signaling between the two cell types.

### SPARC outperforms CAR when both incorporate 4-1BB signaling

To combine the phase separation-mediated signaling enhancement conferred by the NCR1 ICD with the costimulatory signals provided by the 4-1BB, we incorporated both the NCR1 and 4-1BB ICDs into the SPARC, and used two CARs containing both ICDs (NCR1BBz) as comparisons, one carrying the CD8 transmembrane domain and the other carrying the NCR1 transmembrane domain with R-A mutation (**Figure 6A**). Flow cytometric analysis revealed that the surface expression level of SPARC was slightly lower than that of the CAR on NK92MI cells (**Figure S3D**). Upon target cell stimulation, SPARC containing both ICDs induced the highest level of NK cell activation, which was superior to that of the SPARC containing only the NCR1 ICD, and also higher than that of the CAR carrying both ICD (**Figure 6B**). Moreover, SPARC constructs bearing multiple intracellular domains demonstrated stronger activation in NK cells than those containing only a single ICD (**Figure S3E and S3F**). As for primary T cells, the activation level of SPARC-T cells was lower than that of CAR-T upon target cell stimulation, as measured by IL-2 secretion (**Figure 6C**). Flow cytometry showed that the surface expression level of the CAR was higher than that of the SPARC on T cells (**Figure S3G**). We speculate that the lower activation level may be attributable to the lower cell-surface expression of SPARC relative to CAR, which may in turn result from competitive binding of endogenous CD3ζ by the TCR, thereby limiting its availability.

**Figure 6.**
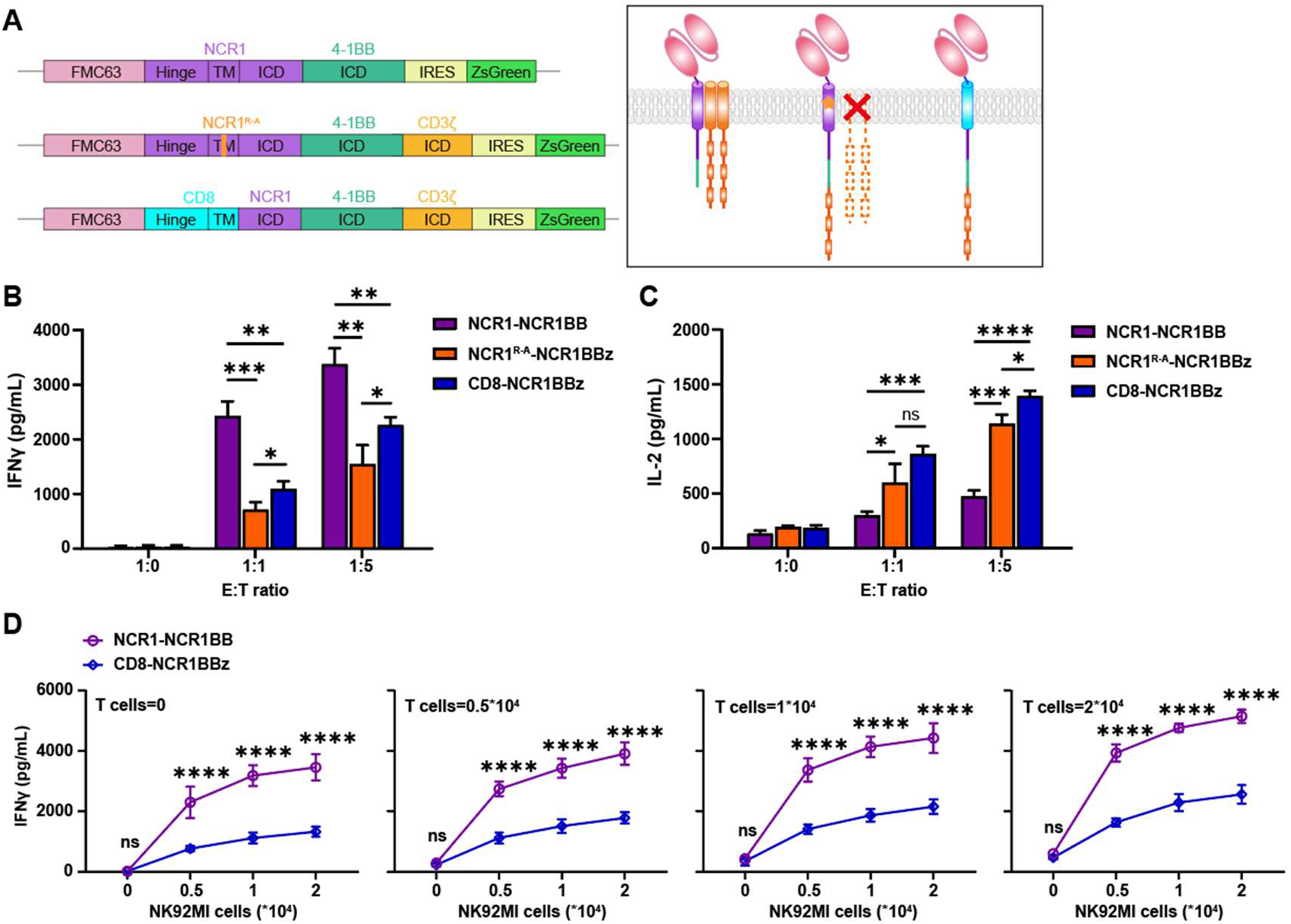
SPARC containing the NCR1 ICD and 4-1BB ICD outperform CAR constructs. **A** Gene (left) and protein (right) schematics of the SPARC with NCR1 and 4-1BB dual ICD, NCR1^R-A^-NCR1BBz and CD8-NCR1BBz CAR. **B** IFN-γ productions of SPARC-NK92MI cells upon Nalm6 cells stimulation for 24 hours. **C** IL-2 productions of SPARC-T cells upon Raji cells stimulation for 24 hours. **D** IFN-γ secretion in the supernatants from co-culture of SPARC- and CAR-NK92MI cells and T cells upon Raji cells stimulation for 24 hours at the indicated cell numbers. Raji cells is 20000 per well. Data are presented as mean with SD (n = 3; ∗, P < 0.05; ∗∗, P < 0.01; ∗∗∗p < 0.001; ∗∗∗∗p < 0.0001; ns, nonsignificant; unpaired two-tailed Student’s t-test for comparisons).

We then engineered both T cells and NK cells with the dual-ICD SPARC or the dual-ICD CARs, and co-cultured them with CD19-expressing Raji lymphoma cells at graded NK:T cell ratios. The results of co-culture experiment demonstrated that when NK and T cells were used in combination, the dual-ICD SPARC elicited markedly higher IFN-γ secretion from both cell types than the NCR1BBz CAR, highlighting their superior application potential (**Figure 6D**). Under conditions with varying T-cell numbers, we fitted curves for IFN-γ secretion in the co-culture supernatant as a function of NK-cell number (**Figure S4A and S4B**). The NK-cell number required to reach half-maximal IFN-γ concentration was lower for dual-ICD SPARC than for dual-ICD CAR, further indicating that dual-ICD SPARC more potently induces cytokine secretion upon NK-cell activation (**Figure S4C**).

## Discussion

In this study, we designed, validated, and characterized a novel class of synthetic parallel antigen receptor complexes (SPARC) that overcome fundamental limitations of conventional CAR therapies by recapitulating the parallel signaling architecture and biophysical mechanisms of natural immune receptors. Our findings demonstrate that this biomimetic approach not only resolves issues of tonic signaling and limited sensitivity but also introduces a new dimension of tunability and synergistic potential for adoptive cell therapy.

The core innovation of the SPARC platform lies in its departure from the tandem, single-chain CAR paradigm. By leveraging the native transmembrane domain of NCR1, KR3DS1 and TCRα to recruit endogenous adaptor proteins (CD3ζ/FcRγ/Dap12), we reconstitute a natural parallel signaling complex. This design inherently avoids the non-specific signaling and steric hindrance associated with artificially fused intracellular domains in traditional CARs.^53^ This novel immunocyte engineering strategy, based on the parallel architecture of receptor complexes, has been proven to be effective and practicable. This architecture permits multiple intracellular domains to exhibit differential signaling dynamics—because they are distributed across distinct protein subunits, each of which precisely recapitulates the native topology of the endogenous receptor.^54, 55^ In contrast, this characteristic is unattainable in conventional single-chain CAR designs. Our data confirm that this electrostatic recruitment is indispensable for robust antigen-specific activation, as mutations disrupting this interaction (R→A or CD8-TM replacement) minimized signaling despite preserved surface expression.

A pivotal and unexpected discovery was the critical role of the NCR1 cytoplasmic domain in signal amplification. Despite its lack of canonical signaling motifs, this short tail was sufficient to confer low-level activation even in the absence of adaptor recruitment. We identified that this function is mediated by two conserved basic-rich sequences (BRS) that drive liquid-liquid phase separation with key signaling kinases such as Lck. The association between BRS and Lck appears to be predominantly governed by electrostatic interactions between oppositely charged residues.^56^ This mechanism facilitates the formation of membrane-proximal condensates that enrich receptors and signaling molecules, thereby enhancing clustering and sensitivity to low antigen density. The abolition of this amplification upon BRS mutation, coupled with direct visualization of receptor clustering via TIRF microscopy, provides compelling evidence that phase separation is a fundamental mechanism underpinning the sensitivity of native NCR1 and, by extension, our SPARC platform. This finding significantly advances our understanding of NK cell biology and establishes phase separation as a powerful design principle for next-generation synthetic receptors. The participation of phase separation in signal transduction has also been documented in the context of transmembrane signaling by other receptor molecules, where it serves as a critical regulator of signal transduction.^41, 57^ Phase-separation modules have also been incorporated into CAR design to improve CAR T cell efficacy.^58, 59^

The modularity of the SPARC system offers a versatile framework for tuning immunotherapeutic responses. We demonstrated that while the NCR1 transmembrane domain is optimal, other natural transmembrane domains (e.g., from TCRα) can be functionally incorporated, albeit with reduced efficacy. This highlights the specificity of natural protein-protein interactions but confirms the generalizability of the parallel assembly concept. Furthermore, the interchangeability of intracellular domains, such as the incorporation of the 4-1BB costimulatory domain, provides a pathway to fine-tune signaling strength and persistence.^60^ This modularity represents a significant advantage over fixed-format CARs, as it enables systematic engineering of receptors with calibrated activation thresholds to maximize tumor killing while minimizing on-target/off-tumor toxicity for diverse clinical contexts. Beyond 4-1BB, our SPARCs also have the potential to provide multiple costimulatory signals by incorporating diverse domains, including OX40, 2B4, and more.^61, 62^

Finally, the translational potential of this platform was underscored by its efficacy in a combined T/NK cell therapy setting. The successful deployment of SPARCs in primary human T cells, which also express the requisite endogenous adaptors, demonstrates the broad applicability of this technology across lymphocyte lineages. The synergistic enhancement of tumor killing observed when SPARC-T and SPARC-NK cells were co-cultured presents a promising strategy to overcome tumor heterogeneity. We postulate that this synergy arises from bidirectional crosstalk, wherein cytokines and membrane-bound ligands mutually activate both cell types, potentially further augmented by the phase-separation capability of the NCR1 tail creating more efficient immune synapses. This combined approach could address key limitations of single-cell-type therapies, such as poor infiltration, antigen escape, and transient persistence.^63–65^

In conclusion, our work establishes a robust and biomimetic framework for synthetic immune receptor engineering. By mimicking the natural parallel architecture of receptors such as NCR1 and harnessing biophysical mechanisms like phase separation, we have established a SPARC platform that effectively elicits antigen-specific immune responses in both NK and T cells. Its inherent modularity and demonstrated synergy in combined T/NK cell therapy pave the way for a new generation of tunable, potent, and persistent immunotherapies for cancer.

## Data Availability Statement

All materials generated or analyzed in this study are included in this published article and its Supplementary Information document.

## Acknowledgement

This work was supported by grants from the National Natural Science Foundation of China (T2394512 and 32200549 to H. C.), the Strategic Priority Research Program of the Chinese Academy of Sciences (XDB1000103 to J. L.), Beijing Medical Award Foundation (YZTZ-2022-0080-0015 to C. L.), and Beijing High Innovation Plan (202504841084 to J. L.). We thank Prof. Xi Wang for helping in NK cell culturing. We thank Dr. Yun Feng and Qing Bian for the help in TIRF-SIM (Total Internal Reflection Fluorescence-Structured Illumination Microscopy) imaging and data processing with Imaris. All the imaging experiments were performed at the Center for Biological Imaging, Institute of Biophysics, Chinese Academy of Science. We thank J. Jia and S. Meng (Core Facility, Institute of Biophysics, CAS) for technical support in the flow cytometry analysis.

## Author contribution

J. L, C. L., and H. C. conceived the project and designed the experiments. Q. W. performed the experiments. S. W. and Y. Q. helped with the experiments. Q. W. prepared the initial figures and drafted the manuscript. All authors contributed to revising the manuscript.

## MATERIALS AND METHODS

### Commercial Antibodies and Protein

Antibodies and protein used in this study were purchased from various commercial suppliers: Anti-human CD69-APC (SinoBiological, 11150-MM06-A); Anti-mouse Fab2-Alexa647 (Jackson Immunoresearch, 115-606-072); Anti-human CD107a-PE (Invitrogen, 12-1079-42); Biotinylated Human CD19 (ACROBiosystems, CD9-H82E9).

### Cell Line and Culture

Human cancer cell lines Jurkat, Nalm6 and Raji were obtained from ATCC. NK92MI cell line was a gift from Dr. Wei Chen (Zhejiang University). Jurkat, Nalm6 and Raji were cultured in RPMI 1640 medium with 10% FBS, 2 mmol/L L-glutamine, and 0.1% penicillin-streptomycin. 293FT cells were cultured in DMEM medium with 10% FBS and 0.1% penicillin-streptomycin. NK92MI and transduced NK92MI were cultured in MEMα supplemented with 0.2 mM inositol, 0.02 mM folic acid, 0.01 mM 2-mercaptoethanol, 0.1% penicillin-streptomycin, 12.5% FBS and 12.5% horse serum. All cells were maintained in 5% CO_2_ at 37 °C.

### Peptides and proteins

#### Synthesized peptides

All peptides used in this study were synthesized by GL biotech (Shanghai). Fluorescence dye labeling was conducted as previously described (Chen et al., 2023).

#### Lck protein

The expression and purification of Lck proteins were conducted as previously described (Chen et al., 2023). Briefly, Lck_UD-SH3-SH2_ (residues 3-226) were expressed in BL21 (DE3) E. Coli (Tsingke) and then purified with His-Trap HP column followed by gel filtration (Hiload 16/600, Superdex 200, Cytia).

### Generation of Lentiviral Constructs, Virus Supernatant, and CAR-NK Cells

Lentivirus constructs encoding CAR and SPARC were generated and cloned into lentiviral transfer plasmid backbone pHAGE with zsGreen reporter linked by IRES. The extracellular domain of CAR construct was generated using the FMC63 scFv fused to the CD8α transmembrane domain, 4-1BB or CD28 intracellular domain and the CD3ζ intracellular domain. The SPARC construct consists of an extracellular FMC63 scFv fused to a specified hinge region, followed by a transmembrane domain and an intracellular domain. NCR1 hinge region, transmembrane domain and intracellular domain (UniprotKB-O76036, amino acids 213-304), NCR3 hinge region, transmembrane domain and intracellular domain (UniprotKB-O14931, amino acids 129-201), KIR3DS1 hinge region, transmembrane domain and intracellular domain (UniprotKB-Q14943, amino acids 316-382), CD16α hinge region, transmembrane domain and intracellular domain (UniprotKB-P08637, amino acids 193-254), TCRα transmembrane domain (UniprotKB-P01848, amino acids 107-139) with K-A/I mutations.

Lentiviral constructs were co-transfected into HEK293FT cells with helper plasmids pMD2.G and psPAX2 using Polyplus transfection reagent (Polyplus, #101000046). Virus-containing culture medium supernatant was collected at 48 and 72 h post transfection. NK92MI and Jurkat cells were transduced with lentiviral supernatants for 24 hours and then cultured in fresh medium. Transduction efficiency was measured 48 to 72 hours post-transduction by flow cytometry.

### Preparation of Engineered Primary T Cells

Lentiviral particles encoding CAR constructs were collected and concentrated, and the resulting viral preparation was applied to target cells in the presence of 8 µg/mL Polybrene. The cells were then spinoculated at 1,600 rpm for 60 min. Surface display of CAR was examined by flow cytometry after staining with anti-mouse Fab-Alexa647 (Jackson ImmunoResearch, #115-606-072).

For the generation of primary engineered T cells, pan CD3+ T cells were purified from PBMCs (Milestone Biotechnologies) using the MojoSort Human CD3 T Cell Isolation Kit (BioLegend, #480022). T cells were activated on day 0 with Human T-Activator CD3/CD28 Dynabeads (Thermo Fisher Scientific, #11161D), transduced with lentivirus on day 1, and then expanded by doubling the medium volume and supplementing with human IL-2 at 2–3-day intervals. On day 7, the Dynabeads were depleted by magnetic separation. Engineered T cells were harvested, and transduction efficiency was determined based on ZsGreen detection and surface receptors staining.

### Flow Cytometry

Cells were harvested for flow cytometry, washed with PBS, pelleted and resuspended in antibody solution, mixed, and incubated for 30 minutes at 4 °C protected from light for surface staining. CAR and SPARC expression on NK92MI and Jurkat cells were measured using a conjugated anti-Mouse Fab-Alexa647 (Jackson Immunoresearch, #115-606-072). The flow data were acquired on BD Influx (BD Biosciences), and data were analyzed using FlowJo software.

### Coculture assay of NK92MI cells and target cells

2×10^4^ NK92MI cells transduced with CAR or SNC were co-cultured with Nalm6 or Raji cells at an effector to target ratio (E:T ratio) of 1:1 – 1:5 in 96-well plate. Cells were co-cultured at 37 °C under 5% atmospheric CO2 for 12-24h. After coculture, plates were centrifuged at 1800 rpm for 5 min. Cells pellets were used for measurements of cytotoxicity, and supernatants were used for measurements of cytokine release.

### Cytotoxicity assays

The cell pellets were resuspended in Annexin-V and PI, mixed, and incubated for 30 minutes at 4 °C protected from light for staining. Cytotoxicity is detected by measuring the apoptosis rate of target cells using flow cytometry.

### Detection of cytokine release by ELISA assay

Cytokine production was measured through enzyme-linked immunosorbent assay (ELISA) using supernatants collected from coculture assays. The cytokine concentrations were measured using following ELISA kits according to the manufacturer’s protocol: human IL-2 (Boster, EK0397), human IFN-γ (Boster, EK0373) and human TNF-α (Boster, EK0525).

### Statistics

All statistical analyses were performed by using GraphPad Prism (Prism 9; GraphPad Software Inc.). The data were analyzed by either one-way ANOVA (multiple groups) or a t-test (two groups) to com pare quantitative differences. Differences with P < 0.05 (two-sided) were considered statistically significant and reported as *, P < 0.05; **, P < 0.01; ***, P < 0.001, ****, P < 0.0001. The data were represented as mean values ± SD, unless mentioned otherwise. The significance test used and sample sizes (n) are reported in each figure legend.

### Preparation of small unilamellar vesicles

A phospholipid mixture comprising 95% DOPC, 5% DGS-NTA-Ni²⁺, and 0.1% DSPE-PEG5000 was evaporated to dryness under a gentle nitrogen stream in a 37 °C water bath. Residual solvent was removed by placing the dried film in a desiccator for 2 h. The film was then hydrated with PBS to a final lipid concentration of 2 mg/mL. Bath sonication was applied for 5–10 min, after which repeated freeze–thaw cycles were performed until the suspension became optically clear. The preparation was centrifuged at 21,380 × g for 60 min at 4 °C, and the supernatant containing small unilamellar vesicles (SUVs) was retained.

### Formation of supported lipid bilayers

Glass-bottomed 96-well plates were cleaned with 5% Hellmanex III and rinsed three times with Milli-Q water. The wells were subsequently treated with 6 M NaOH at 50 °C for 2 h, rinsed extensively with Milli-Q water, and equilibrated with PBS for 30 min. Freshly prepared SUV suspension (15 µL) was added to each well and incubated at 37 °C for 1 h to promote supported lipid bilayer (SLB) assembly. The resulting bilayers were washed three times with clustering buffer (PBS containing 1% BSA) and incubated for 30 min at room temperature.

### Membrane-associated protein condensation assay

His-tagged Lck_UD-SH3-SH2_ (2 µM) was incubated with preformed SLBs for 1 h at room temperature. Unbound protein was removed by three washes with clustering buffer. After an additional 30-min incubation, 10 µM NCR1 cytoplasmic tail or its BRS mutants was introduced into the glass wells and mixed thoroughly to initiate cluster formation. ImageJ was used to measure the occupied area and Feret’s diameters of the resulting clusters. Unless otherwise specified, all imaging was performed on an Olympus FV1200 microscope equipped with a 60× oil-immersion objective.

