## Supplementary Material for "A biomimetic SPARC platform coupling adaptor recruitment and BRS-Lck condensation for enhanced NK/T cell combination therapy"

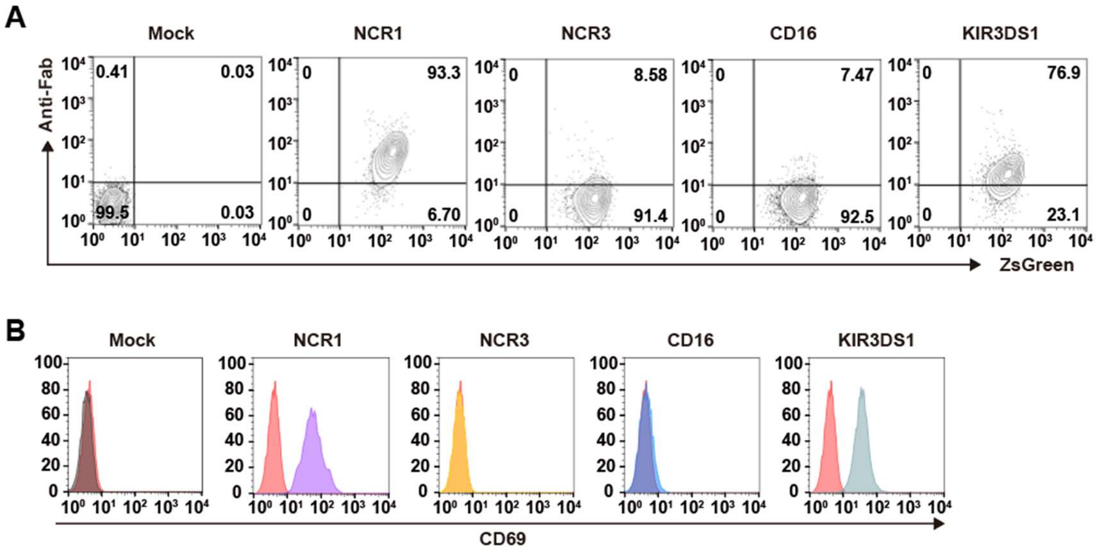


**Figure S1**. Chimeric NK receptor-mediated activation of NK92MI cells.

(**A**) Membrane surface expression of chimeric receptors on NK92MI cell line determined by flow cytometry.

(**B**) Representative data of CD69 expression on transduced NK92MI cells, determined by flow cytometry, after 24 hours coculture with $CD19^+$ Nalm6 cells.

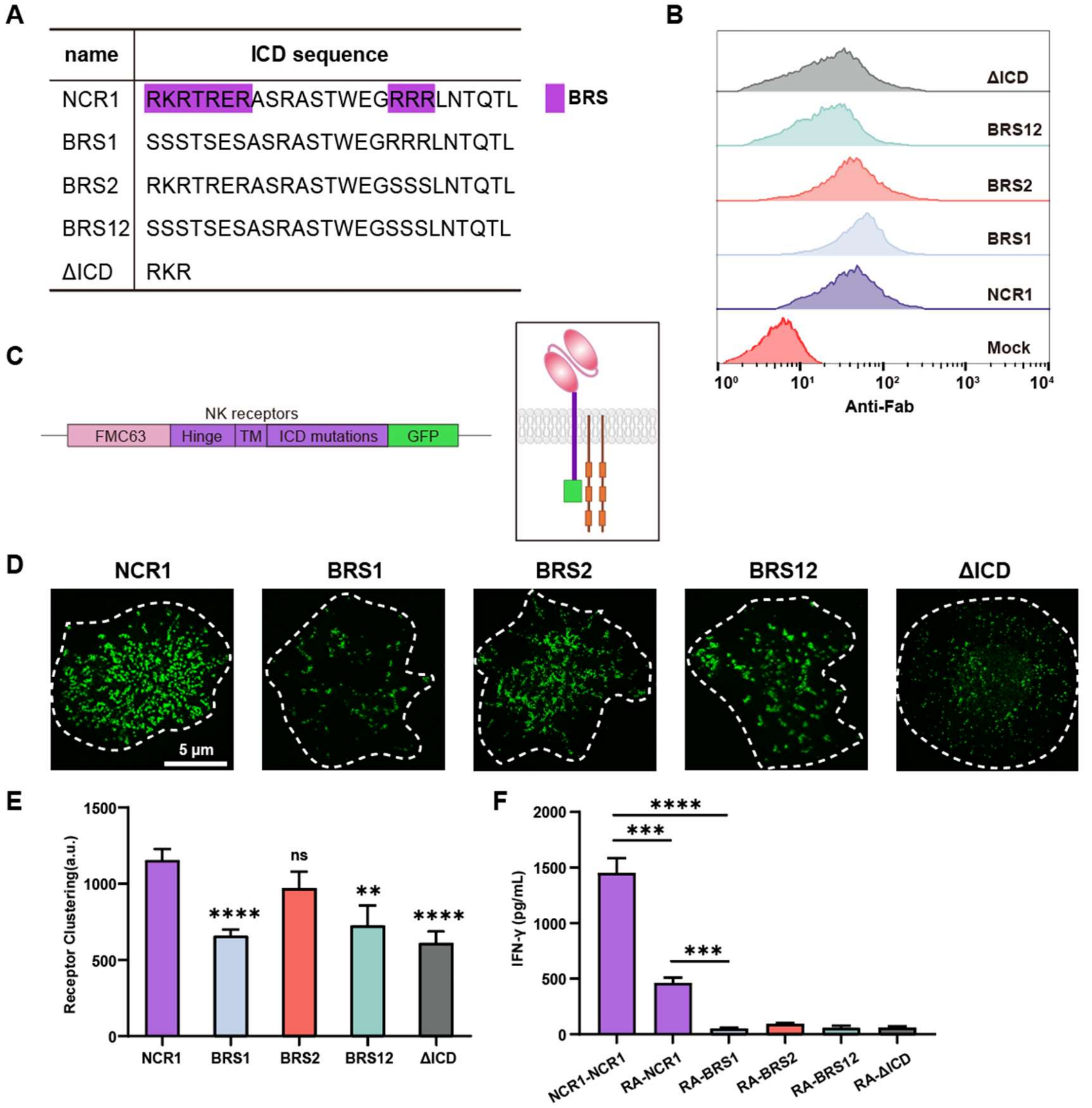


**Figure S2**. NCR1 intracellular domain enhances transmembrane signal transduction via phase separation**.**

(**A**) Amino acid sequences of the NCR1 intracellular domain and its BRS mutants.

(**B**) Surface expression of SPARCs with BRS mutants in (**A**) by flow cytometry.

(**C**) Gene (left) and protein (right) schematics of the SPARCs with GFP tail.

(**D**) Representative results of SPARCs clustering in NK92MI cells stimulated by surface-coated CD19. Fixed NK92MI cells were imaged by TIRF-SIM.

(**E**) Receptors clustering in (**D**) was quantified as described in Materials and Methods (n=20).

(**F**) IFN-γ secretion of SPARC-NK92MI cells with mutations at both domains (TM and ICD) upon Nalm6 cells stimulation for 24 hours at an effector:target (E:T) ratio of 1:5 (n=3).

Data are presented as mean with SD (∗∗p < 0.01; ∗∗∗p < 0.001; ∗∗∗∗p < 0.0001; ns, nonsignificant; unpaired two-tailed Student's t-test for comparisons).

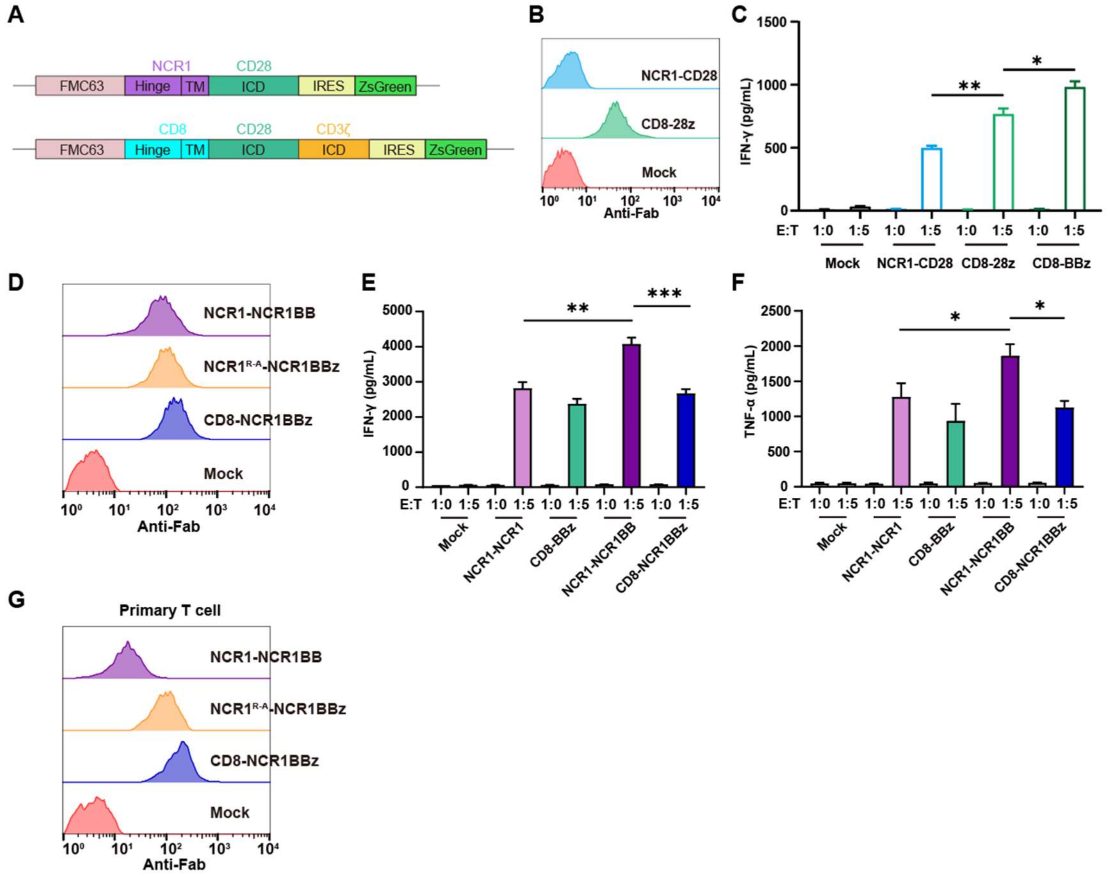

**Figure S3**. Introducing co-stimulatory domain into SPARC achieves better immune efficacy than that of CAR.

(**A**) Gene (left) and protein (right) schematics of the SPARC with CD28 co-stimulatory domain and 28z CAR.

(**B**) Surface expression of SPARC and CAR with CD28 ICD on NK92MI cells by flow cytometry.

(**C**) IFN-γ production by SPARC- and CAR-NK92MI cells after 24 hours stimulation with Nalm6 cells at the indicated effector:target (E:T) ratios.

(**D**) Surface expression of SPARC with NCR1 and 4-1BB dual ICD, NCR1$^{R\text{-}A}$-NCR1BBz and CD8-NCR1BBz CAR on NK92MI cells by flow cytometry.

(**E**) and (**F**) Cytokine productions of SPARC-NK92MI cells upon Nalm6 cells stimulation for 24 hours. IFN-γ (**E**) and TNF-α (**F**) were measured.

(**G**) Surface expression of SPARC with NCR1 and 4-1BB dual ICD, NCR1$^{R\text{-}A}$-NCR1BBz and CD8-NCR1BBz CAR on primary T cells by flow cytometry.

Data are presented as mean with SD (*$p < 0.05$; **$p < 0.01$; unpaired two-tailed Student's t-test for comparisons).

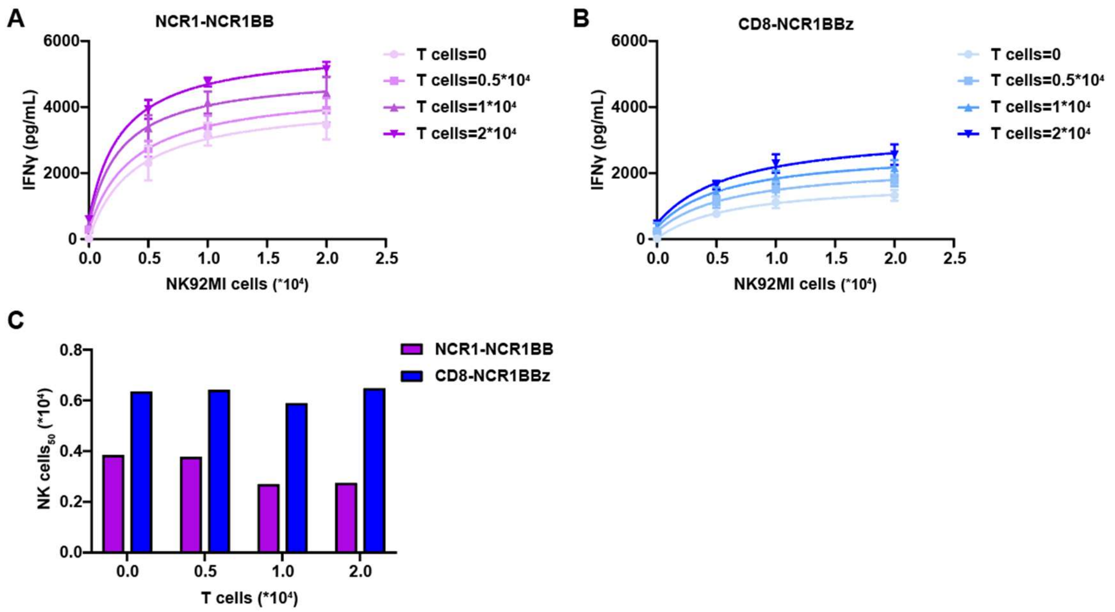


**Figure S4**. SPARC containing the NCR1 ICD and 4-1BB ICD secrete higher levels of IFNγ than CAR constructs.

(**A and B**) Fitted curves of IFN-γ secretion in co-culture supernatants versus NK-cell number for T and NK cells engineered with NCR1-NCR1BB (left) or CD8-NCR1BBz (right).

(**C**) NK-cell number required to reach half-maximal IFN-γ concentration, as derived from the fitted curves above.
